# The effect of biohanin A isolated from the callus culture of meadow clover (*Trifolium pratense L*.) on the life expectancy and survival of *Caenorhabditis elegans*

**DOI:** 10.1101/2022.04.12.488102

**Authors:** Irina Sergeevna Milentyeva, Margarita Yuryevna Drozdova, Violeta Mironovna Le

## Abstract

Oxidative stress, in which healthy aging is limited, is a problem, since in the process there is an uncontrolled production of reactive radicals that negatively affect cells. Heat stress is closely related to oxidative stress, as it stimulates the production of radicals. Meadow clover (*Trifolium pratense L*.) is a promising source of isoflavonoids that have various positive effects on the body. The aim of the work is to study the effect of biohanin A, isolated from *Trifolium pratense L*. callus cultures, on the life expectancy of the *C. elegans* model organism, as well as its survival under oxidative and thermal stress.

It was found that biohanin A increased the lifespan of *C. elegans* worms. The best concentration was determined - 100 microns, at which the survival rate of nematodes after 61 days of incubation was 16.0%. After 48 hours, the survival rate of worms was the highest (87.1%) at a concentration of biohanin A of 200 microns. Other concentrations 10, 50, 100 (81,2, 83,0, 82,4 %) they also showed a higher survival rate compared to the control (74.0%). In addition, the survival rate under heat stress of *C. elegans* was higher by 12.0; 8.4; 4.0% compared with the control. Thus, the results show the antioxidant potential of biohanin A from plant material under oxidative and thermal stress. The substance also had a positive effect on the lifespan of *C. elegans*.

## Introduction

Oxidative stress is a mechanism that limits healthy aging. Aging is a natural process in which there is a functional decrease in organs and tissues associated with the occurrence of various disorders. In the course of life, the body generates reactive oxygen species (ROS), which are a byproduct of oxygen metabolism. With their excessive content, apoptosis, necrosis and cell cycle arrest occur in the tissues of living organisms. To maintain equilibrium, ROS in cells is converted by special enzymes, including superoxide dismutase (SOD), glutathione peroxidase (Gpx), catalase (CAT). In the process of aging, this antioxidant response system deteriorates greatly [5]. In addition, mitochondrial DNA (mtDNA) becomes particularly susceptible to damage during aging. It does not have the protection provided by nucleosomes and repair mechanisms for nuclear DNA [7].

Secondary metabolites of plants are excellent antioxidants that have a milder effect on a living organism in contrast to synthetic ones [2]. Plant *Trifolium pratense L*. it is fodder and has various therapeutic properties. It has various polyphenols in its composition, which are involved in protective mechanisms during stress. The main phytochemical compounds represented by isoflavones include biohanin A, formononetin, daidzein, genistein [1]. Isoflavones are used for osteoporosis, hypertension, menopause. Of particular interest is the plant *Trifolium pratense L*. in the form of a callus cell culture. This raw material source makes it possible to obtain secondary metabolites in higher quantities. Cultures do not require large maintenance costs and in addition, the extraction process is facilitated, since the cells in cultures are not as complex as in full-fledged plants [9].

In this work, the influence of biohanin A isolated from the callus culture of *Trifolium pratense L*. was investigated. on a model organism. The study of the effect of biologically active substances (BAS) can be carried out with the help of soil nematodes *Caenorhabditis elegans*. This model has a number of advantages such as relative ease of maintenance and use, short life span. It has preserved metabolic pathways and more than 200 genes responsible for aging [6].

## Objects and methods

- Callus cultures of meadow clover (*Trifolium pratense L*.) were obtained at the Research Institute of Biotechnology, Kemerovo State University, Russia;
- Preparation of plant extract of callus cultures of meadow clover (*Trifolium pratense L*.): 60% ethyl alcohol was used as an extractant (Russia, Kemerovo Pharmaceutical Factory), milled (grade LZM-1M) and sifted callus cultures were extracted (3 g) at a temperature of 70 ° C in a water bath for 5 hours. The amount of alcohol added is 260 ml. The extraction was carried out twice. In order to prevent the evaporation of ethanol, extraction with a reverse refrigerator was used.
- Isolation of biohanin A from the extract of the callus culture of meadow clover (*Trifolium pratense L*.): Preliminary vacuum evaporation was carried out for the preparative isolation of biohanin A from 60% ethanol extract of *Trifolium pratense L*. callus culture. The temperature was maintained no higher than 50 ° C. Then deionized water was poured in. The volume was 1/4 of the original. After that, the mixture was evaporated until a thick precipitate appeared. The precipitate was treated with n-hexane (5 minutes) using ultrasound. The mixture was processed three times and filtered. The solvent was removed from the combined hood by evaporation under vacuum. After evaporation, the precipitate was added to 50 g of silica gel. The dried precipitate was placed in a column (5×6 cm BioRad) and eluted with a solution consisting of petroleum ether and ethanol in the proportion 99: 1, 98:2, 97:3, 95:5, 93:7, 80: 20. Biohanin A was obtained from evaporated eluates.
- Chemicals and reagents: When preparing the working medium for C. elegans, 1 ml of MgSO4 (1 M), 1 ml of 1 M CaCl2 (1 M), 1 ml of cholesterol (5 mg/ml), 25 ml of K3PO4 (1 M) were added to the sterile agar (NGM). The medium was introduced into Petri dishes in the amount of 20 ml and kept for two to three days at room temperature in order to control the absence of bacterial contamination. To prepare a stock solution of biohanin A, it was dissolved in dimethyl sulfoxide (DMSO). The initial concentration of the solution was 10 M. The initial solution was diluted with sterile distilled water and a concentration of biohanin A - 100, 500, 1000, 2000 microns was obtained. 15 ml of solution was added to the well with nematodes to obtain doses of the substance 10, 50, 100, 200 microns. The solutions were stored at a temperature of 4 °C.
- Strain of Caenorhabditis elegans, content: In the study wild-type nematodes *C. elegans* (strain N2 Bristol) is used. Its cultivation was carried out on cups with NGM medium. To feed the nematodes, an E coli OP50 culture was used, which was previously sown on NGM in the form of a square, without affecting the walls of Petri dishes. Incubation of bacteria without nematodes was carried out for a day at 37 oC. Worms were placed in the center of a cup with bacteria either with a sterile loop or with pieces of agar cut with a scalpel from the previous cup with NGM agar. Incubation of worms with bacteria was carried out at 20 oC. Synchronous worms were obtained as described earlier [8]. Nematodes were also cultured in a liquid S-medium of stage L1. Previously, a nocturnal culture of E. coli OP50 was added on Wednesday at a dose of 0.5 mg/ml. Incubation of bacteria and worms was carried out in a 96-well plate (120 μl) for 48 hours at 20 oC. Then 15 μl of 5-fluoro-2-deoxyuredine (1.2 mM FUDR) was added to the wells. After that, the worms were left for 24 hours at 20 oC to obtain nematodes in the L4 stage.
- Life expectancy analysis: Analysis of the effect of different concentrations (0, 10, 50, 100, 200 microns) of biohanin A was carried out in 96-well plates in a liquid S-medium. The experiment was carried out in 6-fold repetition. After every 4-7 days, the surviving and dead nematodes were counted. The experiment was carried out for 61 days.
- Stress resistance analysis: The effect of biohanin A from plant raw materials on the resistance of *C. elegans* worms to oxidative stress was carried out in a liquid medium on tablets. About 15 ml of biohanin A solution was added to the nematodes in various doses. Incubation of worms with biohanin A in a tablet was carried out at a temperature of 20 C for 5 days. The live and dead *C. elegans* were counted in each cell. To induce oxidative stress, 15 μl of paraquat (1M) was introduced into the cells and the tablet was incubated in a thermostat at a temperature of 20 ° C. After 24 h and 48 h, live and dead *C. elegans* worms were counted. The experimental results obtained were compared with the data for the control group. Control groups of worms were grown without the addition of biohanin A. The effect of biohanin A on the resistance of *C. elegans* worms to temperature stress was carried out in tablets with the addition of a liquid medium. After applying 15 ml of biohanin A solution in various concentrations, the nematodes were kept at a temperature of 20 °C for 5 days. After counting the living and dead C. elegans in each cell, they were transferred to incubation at 33 °C to create thermal stress. After 24 hours and 48 hours, the living and dead *C. elegans* were counted. Control *C. elegans* were grown without the addition of biohanin A.
- Statistical analysis: All experiments were performed in six independent trials. The analysis of statistical data was carried out using the Microsoft Office Excel 2007 software product. The statistical analysis of the obtained data was carried out using the Student’s simultaneous paired criterion, for each pair of interests. The differences were considered statistically significant at p < 0.05.

## Results and discussions

The IR spectrum of isolated and purified biohanin A from callus cultures of meadow clover (*Trifolium pratense L*.) is shown in Figure 1. The degree of purification is at least 95%.

**Figure 1.**
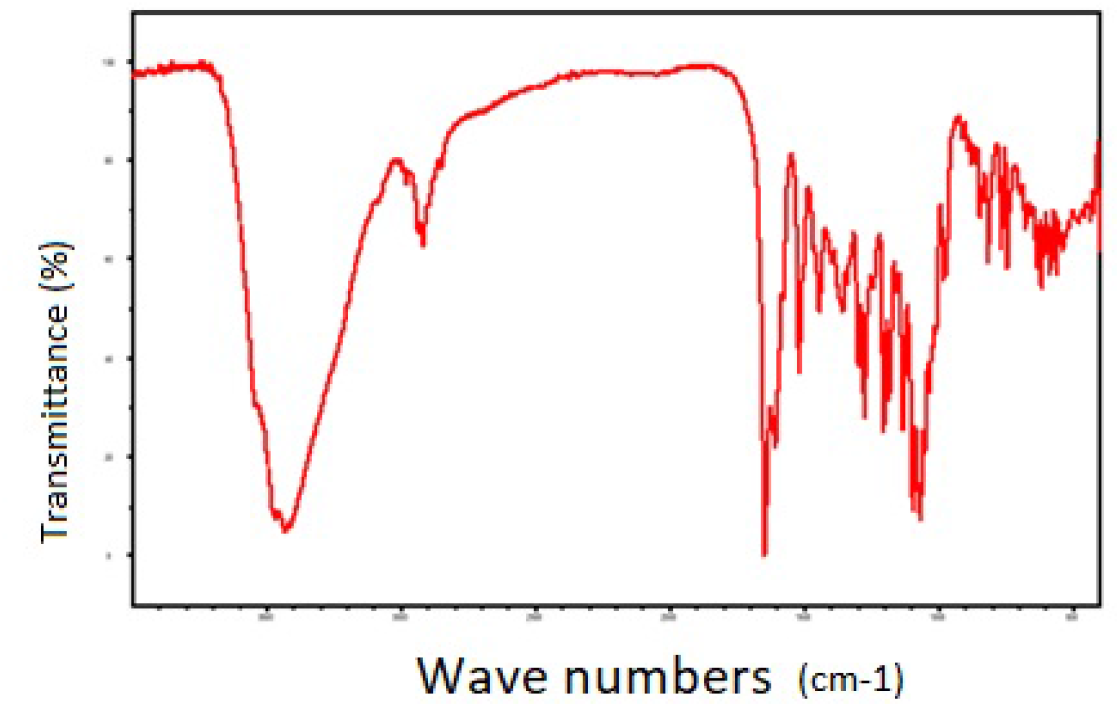
IR spectrum of biohanin A

The *C. elegans* model has proven itself well in testing BAS for life expectancy. So, there are previous studies on the positive effect on longevity of substances such as quercetin [13], boravinone B [8], rutin [11], silymarin [12].

The analysis of life expectancy was carried out using four concentrations of biohanin A – 10, 50, 100, 200 microns (Figure 2). No dose-dependent effect was observed in the experiment. A significant increase in average life expectancy was observed when *C. elegans* worms were treated with biohanin A at a dose of 100 microns. The survival rate of nematodes after 61 days of incubation was higher (16.0%) compared to the control (0.7%). Insignificant survival (3.2 and 2.3%) was shown by other concentrations of BAS of 10 and 50 microns, respectively, compared with the concentration of 100 microns. The concentration of biohanin A – 200 microns showed a negative result on the lifespan of *C. elegans* almost throughout the entire day of incubation of worms. According to our data, this is the first study showing the positive effect of biohanin A isolated from Trifolium pratense L callus cultures on the lifespan of *C. elegans*.

**Figure 2.**
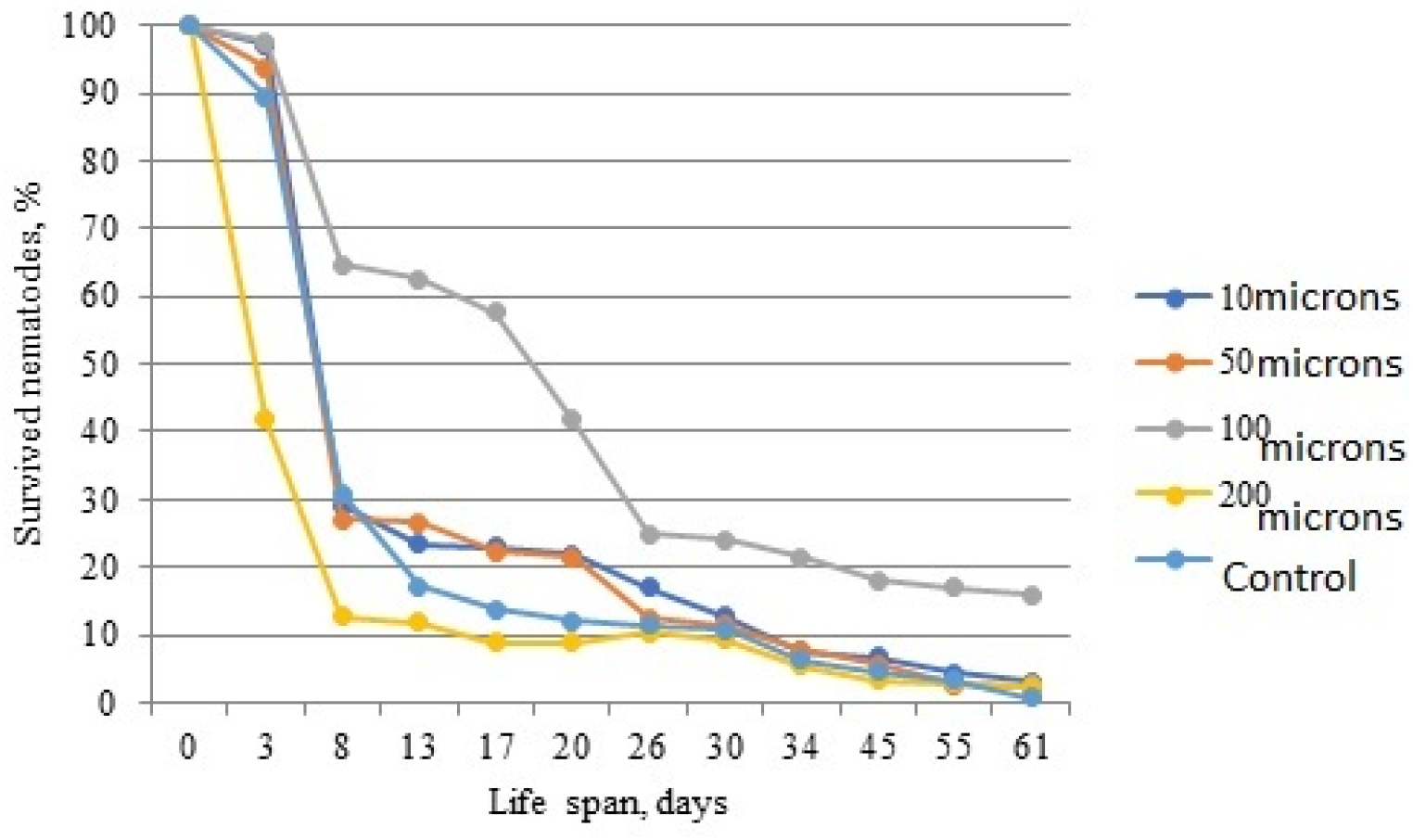
The effect of biohanin A isolated from Trifolium pratense L. callus cultures. for the duration of wild-type worms

There is a strong relationship between aging and stress, as there is a harmful effect of ROS formed during the life of living beings, in particular *C. elegans* [4].

The effect of various tested concentrations of biohanin A on the survival of nematodes under oxidative stress is shown in Figure 3. The following concentrations were used in the experiment - 10-200 microns. Subsequent exposure to ROS generation was carried out by paraquat. The results showed that after 24 hours of incubation, biohanin A isolated from callus cultures of Trifolium pratense L. it did not show significant results, since the survival rate under the influence of all tested concentrations and the control was approximately the same. After 48 hours, the survival rate of worms was the highest (87.1%) at a concentration of biohanin A of 200 microns. Other concentrations 10, 50, 100 (81,2, 83,0, 82,4 %) they also showed a higher survival rate compared to the control (74.0%).

**Figure 3.**
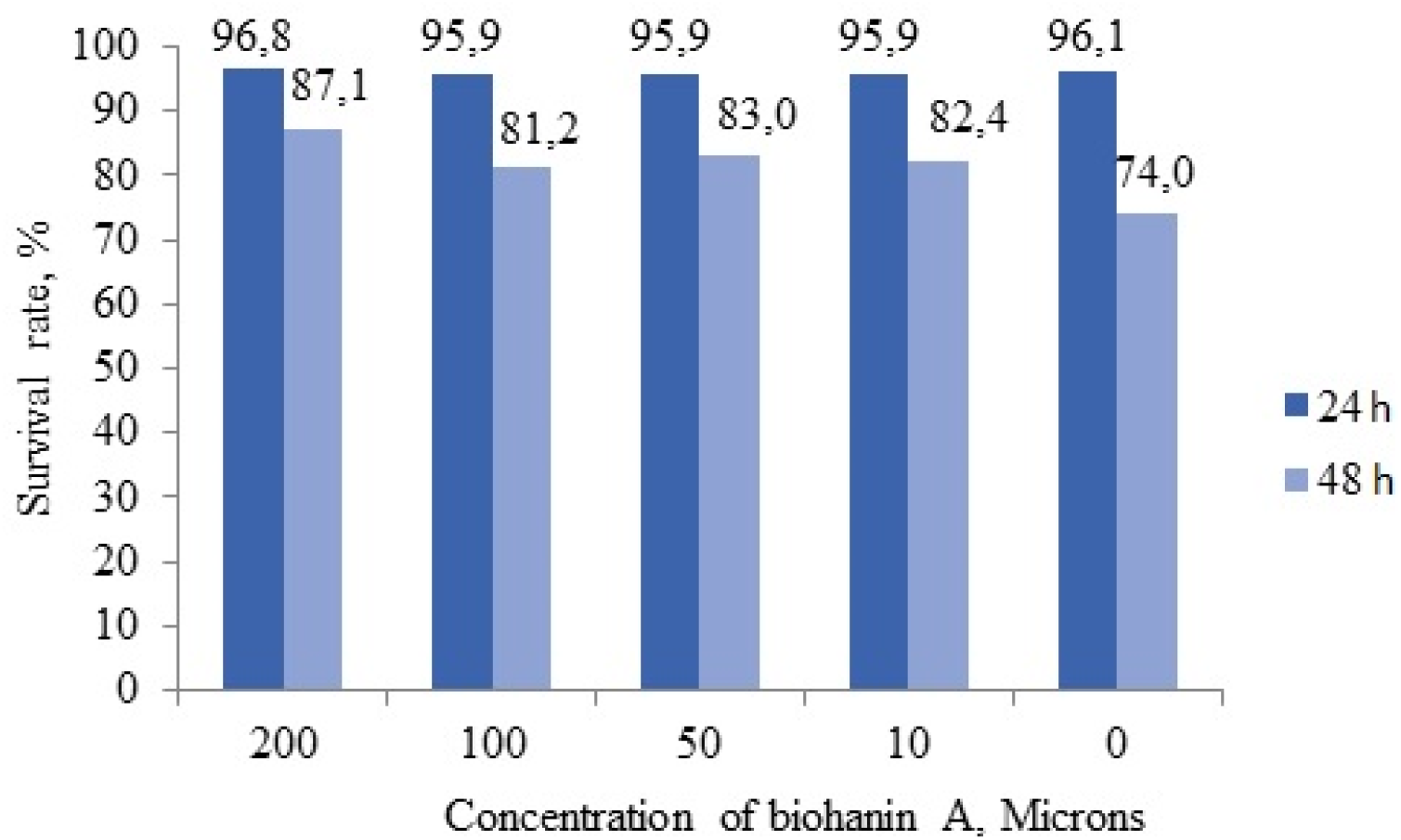
The effect of biohanin A isolated from Trifolium pratense L. callus cultures. on the survival of wild-type worms under oxidative stress

**Figure 3.**
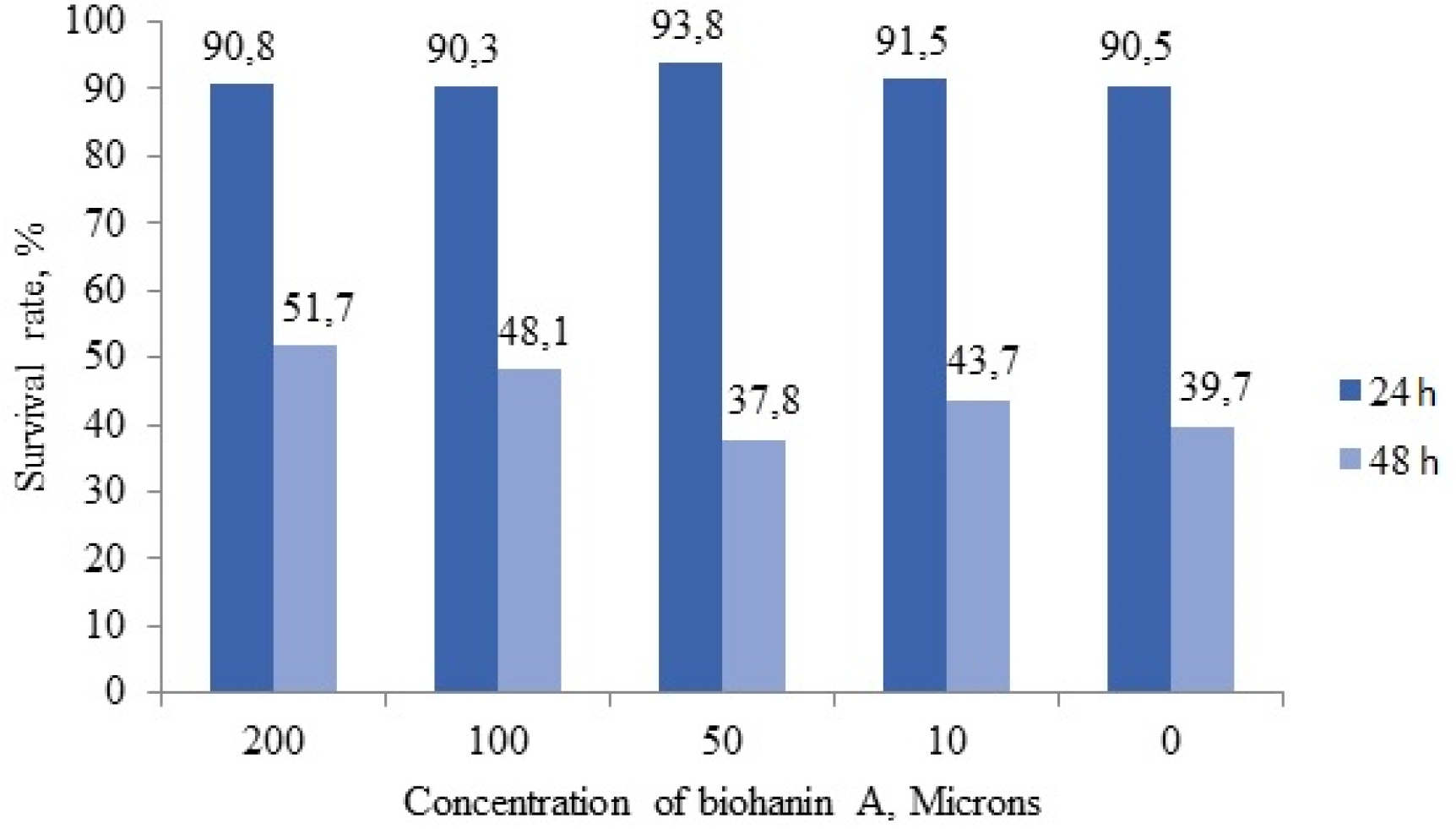
The effect of biohanin A isolated from Trifolium pratense L. callus cultures. on the survival of wild-type worms under heat stress

The oxidative balance can be disrupted by heat stress as a result of high metabolic generation of ROS, as well as reduced activity of the enzymes that eliminate them. In response to various stressful situations, various heat shock proteins can be generated. Thus, under thermal stress, HSF proteins are activated, which affect the suppression of SOD neutralizers, which in turn leads to the accumulation of ROS [10]. In the experiment, after 24 hours of incubation of nematodes under thermal stress, there were no significant differences between the experimental and control groups. After 48 hours, the survival rate of nematodes was higher by 12.0; 8.4; 4.0% compared to the control.

Thus, in the course of the study, a positive effect of biohanin A isolated from callus cultures of meadow clover (*Trifolium pratense L*.) on the model organism of *C. elegans* was revealed in terms of such indicators as life expectancy, survival under stress. It has been shown that biohanin A not only increased the lifespan of worms, but also improved survival under oxidative and thermal stress.

## Notes

### Competing Interest Statement

The authors have declared no competing interest.

